# Genomic Prediction of Strawberry Resistance to Postharvest Fruit Decay Caused by the Fungal Pathogen *Botrytis cinerea*

**DOI:** 10.1101/2021.06.08.447540

**Authors:** Stefan Petrasch, Saskia D. Mesquida-Pesci, Dominique D.A. Pincot, Mitchell J. Feldmann, Cindy M. López, Randi Famula, Michael A. Hardigan, Glenn S. Cole, Steven J. Knapp, Barbara Blanco-Ulate

**Author notes:** Corresponding author: Department of Plant Sciences, University of California, Davis, One Shields Avenue, Davis, California, 95616, USA; and.

## Abstract

Gray mold, a disease of strawberry (*Fragaria* × *ananassa*) caused by the ubiquitous necrotroph *Botrytis cinerea*, renders fruit unmarketable and causes economic losses in the postharvest supply chain. To explore the feasibility of selecting for increased resistance to gray mold, we undertook genetic and genomic prediction studies in strawberry populations segregating for fruit quality and shelf life traits hypothesized to pleiotropically affect susceptibility. As predicted, resistance to gray mold was heritable but quantitative genetically complex. While every individual was susceptible, the speed of symptom progression and severity differed. Narrow-sense heritability ranged from 0.38-0.71 for lesion diameter (LD) and 0.39-0.44 for speed of emergence of external mycelium (EM). Even though significant additive genetic variation was genome wide observed for LD and EM, the phenotypic ranges were comparatively narrow and genome-wide analyses did not identify any large effect loci. Genomic selection accuracy ranged from 0.28-0.59 for LD and 0.37-0.47 for EM. Additive genetic correlations between fruit quality and gray mold resistance traits were consistent with prevailing hypotheses: LD decreased as titratable acidity increased, whereas EM increased as soluble solid whole genome content decreased and firmness increased. We concluded that phenotypic and genomic selection could be regression effective for reducing LD and increasing EM, especially in long shelf life populations, but that a significant fraction of the genetic variation for resistance to gray mold was caused by the pleiotropic effects of fruit quality traits that differ among market and shelf life classes.

## INTRODUCTION

The fleshy fruits produced by strawberry (*Fragaria* × *ananassa*), tomato (*Solanum lycopersicum*), and many other horticulturally important plants are susceptible to postharvest decay by gray mold, a devastating disease caused by the necrotrophic fungal pathogen *Botrytis cinerea* (Jarvis 1962; van Baarlen *et al*. 2007; Williamson *et al*. 2007; Dean *et al*. 2012; Elad *et al*. 2016; Petrasch *et al*. 2019a). *B. cinerea* can infect most organs of the plant but is especially destructive on ripe fruit and senescent tissues of dicotyledonous hosts (Jarvis 1962; Dewey and Grant-Downton 2016). Gray mold renders strawberries unmarketable and often causes significant postharvest losses under conditions favorable for pathogen growth (Barritt 1980; Ries 1995; Dean *et al*. 2012; Petrasch *et al*. 2019a). The mechanisms of defense against *B. cinerea* are physiologically and genetically complex and markedly differ from the gene-for-gene resistance and programmed cell death mechanisms commonly triggered by biotrophic pathogens (Elad and Evensen 1995; Glazebrook 2005; Lorang 2019; Caseys *et al*. 2021). As with other necrotrophic pathogens, *B. cinerea* pathogenesis is promoted by fruit ripening and host cell death (Elad and Evensen 1995; Elad *et al*. 2016; Lorang 2019). Consequently, genetic variation for resistance to gray mold tends to be subtle, limited, and quantitative, which undoubtedly underlies the paucity of studies on breeding for resistance to this pathogen (Finkers *et al*. 2007a; Williamson *et al*. 2007; Rowe and Kliebenstein 2008; Lorang 2019; Zhang *et al*. 2019; Caseys *et al*. 2021).

Because natural genetic resistance has been insufficient to prevent postharvest gray mold disease development, pre-harvest fungicides are often applied to suppress pathogen growth and minimize postharvest losses (Legard *et al*. 1997, 2005; Leroux 2007; Elad *et al*. 2016; Cosseboom *et al*. 2019). Controlling *B. cinerea* with fungicides is difficult because the airborne inoculum is present year round, the host-pathogen interactions are complicated, and the pathogen rapidly evolves resistance to fungicides, particularly after repeated applications of specific chemicals (Jarvis 1962; Leroux 2007; Williamson *et al*. 2007; Cosseboom *et al*. 2019; Zhang *et al*. 2019; Caseys *et al*. 2021). Moreover, pre-harvest foliar applications of fungicides have not been shown to be effective for reducing postharvest gray mold incidence in strawberry fruit possibly because most fruit infections arise from contaminated flower tissues (van Kan 2006; van Baarlen *et al*. 2007; Williamson *et al*. 2007; Veloso and van Kan 2018; Petrasch *et al*. 2019a).

The development of gray mold resistant cultivars has been challenging in strawberry and other hosts because most genotypes are highly susceptible, strong sources of natural genetic resistance have not been identified, and resistance mechanisms are quantitative (Barritt 1980; van Kan 2006; Finkers *et al*. 2007a,b, 2008; Williamson *et al*. 2007; Seijo *et al*. 2008; Lewers *et al*. 2012; Petrasch *et al*. 2019a; Zhang *et al*. 2019; Caseys *et al*. 2021). The feasibility of selecting for increased resistance to gray mold has not been deeply explored in strawberry, a species where limited studies have been undertaken to shed light on the genetics of resistance and assess genetic variation for resistance (Barritt 1980; Seijo *et al*. 2008; Lewers *et al*. 2012). The problem of breeding for resistance to gray mold has been most extensively studied in tomato, albeit without achieving robust or foolproof solutions (Finkers *et al*. 2007a,b, 2008). Genetic studies in tomato and Arabidopsis leaves have identified multiple small-effect quantitative trait loci (QTL) that only account for a small fraction of the genetic variation for resistance, seldom translate across genetic backgrounds, and have not solved the problem of breeding for resistance to gray mold (Finkers *et al*. 2007a,b, 2008; Rowe and Kliebenstein 2008). Although genetic studies of similar depth and breadth have not been undertaken in strawberry, previous studies have not uncovered strong sources of resistance to gray mold (Barritt 1980; Bestfleisch *et al*. 2015; Seijo *et al*. 2008; Lewers *et al*. 2012).

We suspected that selection for increased fruit firmness and other fruit quality traits that extend shelf life pleiotropically increased resistance (decreased susceptibility) to gray mold in strawberry. While hypotheses can be formulated from insights gained from genetic studies in tomato and other hosts (Blanco-Ulate *et al*. 2016a,b; Petrasch *et al*. 2019b; Zhang *et al*. 2019; Caseys *et al*. 2021), natural genetic resistance appears to be negligible and quantitative and additive genetic correlations between gray mold resistance and fruit quality phenotypes are unknown in strawberry (Jarvis 1962; Rhainds *et al*. 2002; Chandler *et al*. 2004; Seijo *et al*. 2008; González *et al*. 2009; Lewers *et al*. 2012; González *et al*. 2013; Bestfleisch *et al*. 2015; Petrasch *et al*. 2019a). The susceptibility of strawberry fruit to *B. cinerea* increases during ripening (Jarvis 1962), which suggests that susceptibility factors accumulate independent of defense mechanisms during fruit maturation and senescence, as is typical for this necrotroph (Williamson *et al*. 2007; Zhang *et al*. 2019; Caseys *et al*. 2021; Silva *et al*. 2021). Changes in fruit firmness and other fruit quality traits associated with fruit maturation and ripening in tomato have been shown to increase susceptibility to *B. cinerea* (Blanco-Ulate *et al*. 2016a,b; Silva *et al*. 2021). Although previous studies have been somewhat inconclusive in strawberry, firm-fruited cultivars are predicted to be more resistant to *B. cinerea* than soft-fruited cultivars (Gooding 1976; Barritt 1980). Moreover, ripening-induced differences in proanthocyanidin and anthocyanin accumulation have been predicted to affect *B. cinerea* resistance in tomato and strawberry (Jersch *et al*. 1989; Zhang *et al*. 2013; Bassolino *et al*. 2013).

To more deeply explore the genetics of resistance to gray mold and assess the feasibility of applying genomic selection for increased resistance to gray mold in strawberry, we developed and studied training populations segregating for fruit quality traits predicted to affect shelf life. Genomic prediction approaches are particularly attractive for postharvest traits that are difficult and costly to phenotype in strawberry but still require sufficient accuracy to complement phenotypic selection and achieve genetic gains (Heffner *et al*. 2010; Jannink *et al*. 2010; Lin *et al*. 2014; VanRaden 2020). The training populations for our studies were developed from crosses between firm-fruited long shelf life (LSL) cultivars and soft-fruited short shelf life (SSL) cultivars. Although the gray mold resistance phenotypes of the parents of these populations were unknown, our hypothesis was that selection for extended shelf life has pleiotropically increased resistance to gray mold in strawberry, primarily because fruit of long shelf life cultivars deteriorate more slowly in postharvest storage than those of short shelf life cultivars. We describe a highly repeatable artificial inoculation protocol for gray mold resistance phenotyping developed for the genomic selection studies described herein. Finally, we discuss the prospects for increasing genetic gains for resistance to gray mold through the application of genomic prediction approaches.

## MATERIALS AND METHODS

### Plant Materials and Study Design: Shelf Life Assessment of Modern Long Shelf Life Cultivars

Shelf life studies were conducted with fruit harvested from five day-neutral cultivars (‘UCD Royal Royce’, ‘UCD Valiant’, ‘UCD Moxie’, ‘Cabrillo’, and ‘Monterey’) and three ‘summer-plant’ (‘UCD Finn’, ‘UCD Mojo’, and ‘Portola’) cultivars (the ‘UCD’ prefixes are hereafter dropped from the cultivar names) grown on organic and conventional farms using the standard production practices of commercial growers in coastal California. The day-neutral cultivars were grown in 20-plant plots on three commercial farms, one organic and two conventional, in Santa Maria and Prunedale, California in 2017-18 with harvests for postharvest studies on June 22 and 27, July 30, and August 1, 2018, 7/30/2018. The summer-plant cultivars were grown in 20-plant plots on three commercial farms, one organic and two conventional, in Oxnard and Santa Maria, CA in 2018-19 with harvests for postharvest studies on September 27 and 30, November 1 and 18, and December 4, 2019.

To assess shelf life and estimate gray mold incidence, fruit was harvested on two dates at each location and stored in a dark walk-in cooler maintained at approximately 4°C and 90-95% relative humidity for 21 days postharvest (dph). Harvest dates were June 22 and 27, July 30, and August 1, 2018 for day-neutral cultivars and September 27 and 30, November 1, 18, and 21, and December 4, 2019 for summer-plant cultivars. Two 0.45 kg samples of fruit were collected at each harvest and placed in 18.4 cm × 12.1 cm × 6.2 cm vented plastic clamshells, one of which was stored undisturbed for visual phenotyping and another of which was used for destructive fruit quality trait measurements at three time points for summer-plant experiment (0, 7, and 14 days postharvest; dph) and four time points for the day neutral experiment (0, 7, 14, and 21 dph). Soluble solid content (SSC =°Brix), fruit firmness (g-force), fruit weight (g/fruit), fungal decay incidence (% of clamshell), and marketability were recorded at each time point. The latter was visually scored on a 1 to 5 scale, where 1 = very good, 2 = good, 3 = fair, 4 = poor, and 5 = very poor (Mitcham *et al*. 1996; do Nascimento Nunes 2015). Four fruit were randomly selected from each clamshell at each time point for fruit firmness and SSC measurements. Firmness was measured using a handheld penetrometer (QA Supplies Model FT02) with a 3 mm probe. Soluble solid content (SSC; °Brix) was measured in the juice of macerated fruit with a digital hand-held refractometer (Atago Model PAL-1). Statistical analyses of these experiments were separately performed using the R package *lme4* (Bates *et al*. 2015) with cultivar as a fixed effect and location, cultivar × location, harvest, cultivar × harvest, harvest × location, and cultivar × location × harvest as random effects. Estimated marginal means (EMMs) and linear contrasts between cultivar EMMs were estimated using the R package *emmeans* (Lenth 2021).

### Plant Materials and Study Design: Genetics of Gray Mold Resistance

Seeds of five *F*. × *ananassa* full-sib families were harvested from crosses produced in a greenhouse at UC Davis in the winter of 2018: Royal Royce × Primella (PI551422), Royal Royce × Madame Moutot (PI551632), Royal Royce × Tangi (PI551481), Royal Royce × Earlimiss (PI551862), and 05C197P002 × 16C108P065. These families constituted the multi-family training population developed for genomic prediction and other analyses. Seeds were scarified and germinated the week of 18-22 June 2018. Seedlings were established and grown in a greenhouse until they were transplanted to the field on 5 October 2018 at the UC Davis Wolfskill Experiment Orchard (WEO), Winters, CA. The field site was prepared with a disk-harrow and ring-roller and smoothed with a spring-tooth harrow. The soil was fumigated on 14 May 2018 with Pic-Clor 60® (1, 3-dichloropropene 39% and chloropicrin 59.6%; Cardinal Professional Products, Woodland, CA) at a rate of 474.7 kg/ha. Subsequent to fumigation, planting beds were mechanically shaped to a height of 30.5 cm, width of 50.8 cm, base of 121.9 cm with center-to-center spacing of 152.4 cm between beds, and a furrow width of 30.5 cm. Seedlings were transplanted to planting beds on 15 October 2018 in a single centered row with 55.9 cm between plants within the row. The number of individuals that produced sufficient fruit for the postharvest study of resistance to gray mold were *n =* 86 for Royal Royce × Primella, *n* = 82 for Royal Royce × Madame Moutot, *n* = 78 for Royal Royce × Tangi, *n* = 92 for Royal Royce × Earlimiss, and *n* = 42 for 05C197P002 × 16C108P065. The parents were planted from bare-root plants produced in a commercial high-elevation nursery in Dorris, CA. The population was grown through 30 June 2019, irrigated as needed to prevent water stress, and hand weeded throughout the growing season. Fruit were harvested on from 15 May to 7 June 2019. Additional Royal Royce × Tangi full-sib individuals (*n* =155) were grown, handled, and phenotyped in 2019-20 exactly as described above for the 2018-19 experiment (phenotypic data were collected for 233 Royal Royce × Tangi individuals over two years). Fruit were harvested 8-29 May 2020.

### DNA Isolation and SNP Marker Genotyping

DNA was extracted from 0.2 g of dried young leaf tissue with the E-Z 96 Plant DNA Kit (Omega Bio-Tek, Norcross, GA) per the manufacturer’s instructions, though the protocol was modified by adding Proteinase K to the lysis buffer to a final concentration of 0.2 mg/ml and extending lysis incubation to 45 min. at 65°C in order to increase the quality and yield of the DNA. Single nucleotide polymorphisms (SNPs) were genotyped using the 50K Axiom SNP array (Hardigan *et al*. 2020), and SNP calls were generated using the Affymetrix Axiom Suite (v1.1.1.66). The raw genotypic data were filtered to identify polymorphic SNP markers with clear and well separated homozygous and heterozygous genotypic classes and eliminate individual SNP markers with minor allele frequencies (MAFs) < 0.05 and any missing data. This process yielded 11,946 SNPs for the training population (*n* =380) studied in 2019 and 9,962 SNPs for the Royal Royce × Tangi full-sib population (*n* = 233) studied in 2019 and 2020.

### Gray Mold Resistance Phenotyping

We developed a high-throughput protocol for postharvest phenotyping of *B. cinerea* disease progression and symptom development on ripe fruit. Spore suspensions of the *B. cinerea* strain B05.10 (Büttner *et al*. 1994; Quidde *et al*. 1998) were produced from spores grown on potato dextrose agar as described by Petrasch *et al*. (2019b). Uniformly ripe fruit were harvested at sunrise, avoiding fruit that were under- or over-ripe. The fruit were immediately transferred to cold storage (2.5°C) and inoculated the day of harvest. Several incubation temperatures (2.5, 5.0, 10.0, and 20.0°C) were tested to identify the optimum temperature for *B. cinerea* growth and development with a minimum of contamination from other postharvest decay pathogens. Fruit were placed on 30-cell plastic egg hatching trays with dimensions of 29 cm × 29 cm and 4.5 cm × 4.5 cm cells. The fruit were punctured once near the center with a 3 mm sterile pipette tip to an approximate depth of 1-2 mm. Ten *μ*l of the *B. cinerea* conidia suspension (500 conidia/*μ*l) was placed on the surface of the puncture. The inoculated fruit were incubated in a growth chamber at 10°C and 95% humidity for 14 days. Disease symptoms were assessed daily after inoculation by manually measuring lesion diameter (LD) and determining the number of days until external mycelium (white or gray hyphae) was evident on the surface of the fruit (EM) near the wound site. Fruit were phenotyped until mycelia covered the entire surface of the fruit. Spoiled fruit with infections outside of the inoculation site or caused by decay organisms other than *B. cinerea* were removed from the experiment. Genome-wide association study (GWAS), QTL mapping, and genomic selection (GS) analyses were applied to LD at 8 days post-inoculation (dpi) and EM.

### Fruit Quality Phenotyping

Fruit quality phenotypes were measured on four fruit harvested from individuals in the multi-family population at harvest. The fruit were photographed with a Sony *α* 6000 camera equipped with an E PZ 16–50 mm F3.5–5.6 OSS lens (SONY, Tokyo, Japan). Photographs were processed with a custom macro in Fiji (Schindelin *et al*. 2012; Rueden *et al*. 2017) to obtain RGB color metrics (File S1). RGB colors were subsequently converted into Lab colors using the convertColor() function in R (R Core Team 2021). Fruit firmness (maximum resistance g-force) and fruit diameter (mm) was assessed on whole fruit using a TA.XT plus Texture Analyzer with a TA-53 3 mm puncture probe (Stable Micro Systems Ltd., Goldaming, United Kingdoms). Fruit samples were frozen at −20 °C in Whirl-Pak® Homogenizer Blender Filter Bags (Nasco, Fort Atkinson, WI, USA) for quantifying titrable acidity (TA; %), soluble solid content (SSC; °BRIX), and total anthocyanin concentration (AC; *μg*/*ml*). TA percentages were quantified with a Metrohm Robotic Titrosampler System from 1-5 ml of the defrosted homogenized fruit juice (Metrohm AG, Herisau, Switzerland). SSC was measured from approximately 200 μl of juice on an RX-5000*α*-Bev Refractometer (ATAGO Co. Ltd., Tokyo, Japan). Total anthocyanin concentration was measured from a 25 μl sample of juice in 200 μl 1% HCl in methanol by reading absorption at a wavelength of 520 nm on a Synergy HTX platereader equipped with Gen5 software (Molecular Devices, San Jose, California, USA). A standard curve (*y* = *sx* + *i*) was calculated for quantifying AC using a dilution series of pelargonidin (Sigma Aldrich, St. Louis, MI, USA) from zero to 300 μg/ml in 50 μg/ml increments, where *y* were absorption readings for the perlagonin dilution series, *s* was the slope, *x* was the concentration of perlagonin in the dilution series, and *i* was the intercept. AC was estimated by (*A* – *i*) /*s*, where *A* was the absorption reading.

### Statistical Analyses: Genetic and QTL Mapping and Genome Wide Association Study

Genome-wide association study (GWAS) approaches were applied to search for marker-trait associations in the multi-family training population and the Royal Royce × Tangi population. Estimated marginal means (EMMs) for LD and EM were estimated from sub-samples and biological replications using the *R* package *emmeans* (Lenth 2021). GWAS analyses were applied using the GWAS() function in the R package *rrBLUP* (Endelman 2011). The genomic relationship matrix was used to correct for population stratification (Endelman 2011). The genomic inflation factor (*λ*) was 0.60 for LD and 0.71 for EM for analyses of the multi-family population and 1.09 for LD and 1.00 for EM for analyses of the Royal Royce × Tangi population. The Bonferroni-corrected threshold for statistical significance was -log_10_(0.05/k) where k is the number of SNPs used in the analysis. The Bonferroni-corrected threshold was -log_10_(4.2 × 10^6^) for the multi-family population and -log_10_(5.0 × 10^6^) for the Royal Royce × Tangi population.

Parent-specific genetic maps were developed for each full-sib family using a custom PERL script pipeline utilizing the R packages *BatchMap* and *onemap* (Schiffthaler *et al*. 2017; Margarido *et al*. 2007) (File S4). This pipeline was used to bin co-segregating markers, calculate pairwise recombination fractions, assign markers to linkage groups, and estimate linkage disequilibrium (LD) statis-tics between groups of markers. Specifically, initial linkage groups (representing chromosomal fragments or sub-linkage groups) were assembled using a LOD threshold of 10.0 and a maximum recombination fraction of 0.08. These thresholds typically generate more linkage groups that chromosomes. These chromosomal fragments or sub-linkage groups were then merged manually based on inter-group linkage disequilibrium statistics and percent-identity against the physical genome (Edger *et al*. 2019). The RECORD algorithm (Van Os *et al*. 2005) was used to estimate marker order and genetic distances across a sliding window of 25 markers with a window overlap of 18 markers.

QTL analyses were applied to parent-specific genetic maps within each full-sib family using the scanone() function and Haley-Knott regression (Haley and Knott 1992) as implemented in the R package *qtl* (Broman *et al*. 2003). The null hypothesis of no significant difference between SNP marker genotypes was tested for each locus using backcross equivalent contrasts; specifically, *ȳ_Aa_* – *ȳ_aa_* for SNP markers segregating in the female parent and *ȳ_aa_* – *ȳ_Aa_* for SNP markers segregating in the male parent, where *Aa* is a heterozygote and *aa* is a homozygote. The null hypothesis of no QTL effect was rejected when the likelihood odds (LOD) ratio for the SNP marker effect exceeded the genome-wide LOD significance threshold empirically estimated by permutation with 1,000 randomly drawn samples (Churchill and Doerge 1994).

We searched the *F*. × *ananassa* ‘Camarosa’ reference genome (Edger *et al*. 2019) for QTL-associated candidate genes with putative biotic stress or disease resistance gene function annotations. Gene Ontology (GO) annotations were filtered to identify candidate genes predicted to be involved in plant-pathogen interactions as described by Silva *et al*. (2020). Candidate genes were annotated using the KEGG Automated Annotation Server (KAAS) (Moriya *et al*. 2007) pipeline and filtered pathways as described by Silva *et al*. (2020). The iTAK pipeline was used to predict presence of transcription factors and protein kinases (Zheng *et al*. 2016).

### Statistical Analyses: Estimation of Genetic and Genomic Prediction Parameters

The repeatability on a progeny mean-basis (*R*) was estimated for each trait from multiple subsamples/individual (fruit/individual) by 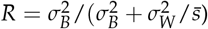, where 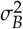 is the between-individual variance component, 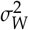 is the among subsamples nested in individuals variance component, 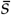 is the harmonic mean number of subsamples/individual, and 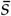 was 2.4 in the multi-family population and 3.6 in the Royal Royce × Tangi population. Variance components were estimated using REML in the R package *lme4* (Bates *et al*. 2015). Narrow-sense genomic heritability (*h*^2^) was estimated for each trait from the mean of 1,000 REML estimates of the additive genetic and phenotypic variance components from 1,000 Markov Chain Monte Carlo (MCMC) samples drawn for cross-validation using G-BLUP (Endelman 2011).

Genomic estimated breeding values (GEBVs) were estimated for each trait by applying three whole-genome regression methods in both training populations—genomic best linear unbiased prediction (G-BLUP), reproducing kernel Hilbert spaces (RKHS), and support vector machine (SVM). G-BLUP, RKHS, and SVM mixed model analyses were performed using the kinBLUP() function of *rrBLUP* (Endelman 2011), the BGLR() function of the R package *BGLR* (Pérez and de Los Campos 2014), and the svm() function of R package *e1071* (Meyer *et al*. 2019), respectively. The kernel for RKHS was determined using the multi-kernel averaging method (de Los Campos *et al*. 2010). Cross-validation analyses were performed for each population × trait × whole genome regression (WGR) method using 1,000 MCMC samples/analysis, where GEBVs were estimated from a random sample of 80% of the individuals and predicted for the GEBVs for the other 20% of the individuals in each sample: 100% of the subsamples/individual were used for these analyses. We replicated these cross-validation analyses using a single randomly selected subsample/individual, where GEBVs were estimated from a random sample of 80% of the individuals (with a single subsample/individual) and predicted the GEBVs for the other 20% of the individuals. Finally, these cross-validation analyses were repeated by randomly selecting a single subsample/individual from 100% of the individuals and estimating the correlation between the single-subsample GEBVs with the EMMs estimated from 100% of the subsamples/individual. Genomic selection accuracy was estimated for each of the nine analyses as the correlation between the phenotypic means (EMMs) and GEBVs (*r_ȳ_*,*GEBV*) for each trait from 1,000 MCMC samples/analysis.

### Data Availability

Genotypic (File S2), phenotypic (File S3), and other supplemental data files, figures, and tables are centrally available at https://figshare.com). File S1 is a text file with a custom macro developed to process photographic images of strawberry fruit using ImageJ, a “public domain Java image processing program” (https://imagej.nih.gov/ij/index.html). File S2 stores 50K Axiom SNP array genotypic data for the multifamily and the Royal Royce × Tangi population. File S3 stores the phenotypic data from our study. Custom R and PERL scripts developed for genetic mapping analyses are stored in File S4. File S5 stores information on candidate genes linked to gray mold resistance QTL with physical addresses and genome annotations from the *F*. × *ananassa* ‘Camarosa’ v1.0 reference genome (Edger *et al*. 2019). The latter is available at the Genome Database for Rosaceae (https://www.rosaceae.org/species/fragaria_x_ananassa/genome_v1.0.a1). File S6 stores the estimated marginal means (EMMs) and genomic-estimated breeding values (GEBVs) for gray mold resistance and fruit quality phenotypes among 380 individuals in a multi-family training population and 233 individuals in the Royal Royce × Tangi population. Fig. S1 displays time-series photographs of fruit of the long shelf life summer-plant cultivars ‘Finn’ and ‘Mojo’ at 0, 7, and 14 days of postharvest storage. Fig. S2 displays the phenotypic distributions and parent EMMs for gray mold resistance phenotypes within full-sib families. Fig. S3 displays kernel densities for genomic prediction accuracy estimated by cross-validation using support vector machine.

## RESULTS AND DISCUSSION

### Natural Postharvest Gray Mold Infections on Fruit of Long Shelf Life Cultivars

Our studies were partly motivated by the observation that gray mold infections were uncommon between 0 and 14 dpi in a series of postharvest shelf life studies of modern long shelf life (LSL) strawberry cultivars (Fig. 1; Fig. S1). The fruit for these studies were harvested from five day-neutral cultivars (‘Royal Royce’, ‘Valiant’, ‘Moxie’, ‘Cabrillo’, and ‘Monterey’) and three summer-plant cultivars (‘Finn’, ‘Mojo’, and ‘Portola’) grown on commercial farms in coastal California (Fig. 1; Fig. S1). These studies produced several insights. Gray mold infections were rarely observed before 14 dph on any of the cultivars tested (Fig.1; Fig. S1). Statistically significant differences in gray mold incidence were not observed among day-neutral cultivars (*p* = 0.87) or summer-plant cultivars (*p* = 0.98). Gray mold incidence ranged from 0.0 to 2.7% among cultivars at 14 dph, a typical postharvest storage duration for LSL cultivars. The five day-neutral cultivars were screened out to 21 dph to gain insights into the postharvest storage limits for modern LSL cultivars (Fig. 1). Although the fruit were still marketable at 14 dph, they became marginally marketable or unmarketable by 17-18 dph (Fig. 1). We observed an exponential increase in gray mold incidence beyond 17-18 dph for every cultivar with means ranging from 10.3 to 36.7% among day-neutral cultivars at 21 dph. These studies showed that gray mold was ubiquitous and eventually rendered the fruit unmarketable but that the natural incidence of gray mold was negligible on fruit of LSL cultivars grown in coastal California within the 14 day postharvest storage window (Fig. 1; File S2).

**Figure 1.**
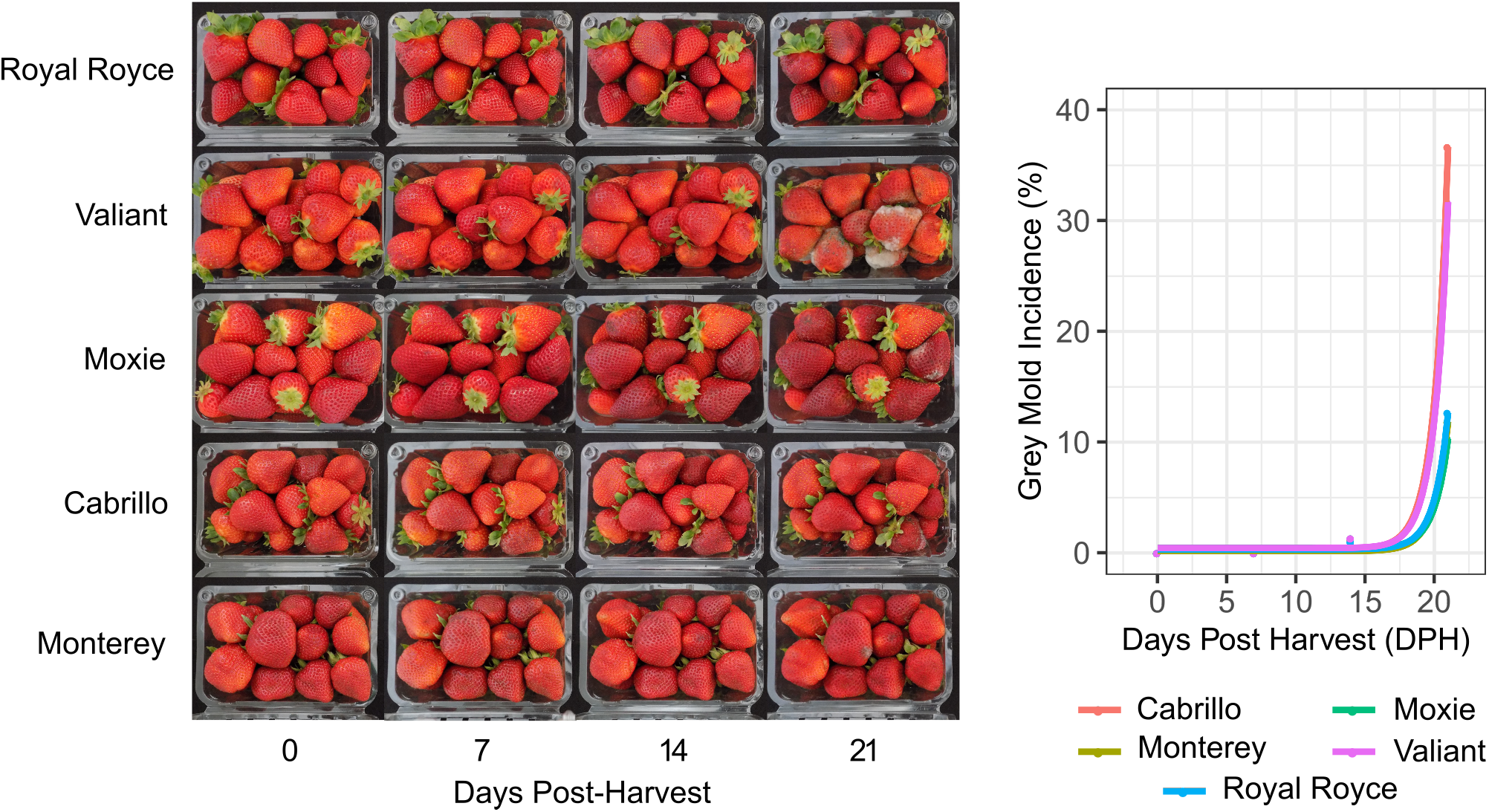
Postharvest Visual Appearance of Cold Stored Fruit of Long Shelf Life Day-Neutral Cultivars Grown in Coastal California. Fruit of ‘Royal Royce’, ‘Valiant’, ‘Moxie’, ‘Cabrillo’, and ‘Monterey’ were harvested June 22 and 27, July 30, and August 1, 2018 from commercial farms in Santa Maria and Prunedale, California, immediately cooled, stored undisturbed in 0.45 kg clamshells at 4°C and 90-95% relative humidity for 21 days postharvest (dph), and photographed and phenotyped 0, 7, 14, and 21 dph (lefthand panel). Estimated marginal means for gray mold incidence (%) were estimated and plotted and exponential regressions were fit to the original phenotypic observations (right panel). *R*^2^ estimates for goodness-of-fit of the exponential functions were 0.59 for Cabrillo, 0.22 for Monterey, 0.25 for Moxie, 0.25 for Royal Royce, and 0.50 for Valiant.

From previous surveys of phenotypic diversity for resistance to gray mold and common knowledge (Gooding 1976; Barritt 1980; Lewers *et al*. 2012), we hypothesized that the low incidence of gray mold on commercially produced fruit of LSL cultivars might be genetically correlated with fruit firmness and other fruit quality traits affecting shelf life (Fig. 2). Although phenotypic correlations have been reported (Barritt 1980), genetic correlations have not. The fruit of LSL cultivars are typically much firmer than the fruit of short shelf life (SSL) cultivars commonly grown for local or direct-market consumption, as exemplified by Earlimiss, Madame Moutot, and Primella in the present study (Fig. 2). The latter are sweeter, softer, and perish more rapidly than ‘Royal Royce’ and other LSL cultivars under normal postharvest storage conditions (Fig. 2). To explore how these phenotypic differences affect resistance to gray mold, we developed a training population (n =380) for genomic selection studies by crossing ‘Royal Royce’, one of the LSL cultivars assessed for natural infections (Fig. 1), with four SSL cultivars (‘Earlimiss’, ‘Madame Moutot’, ‘Primella’, and ‘Tangi’), in addition to crossing a pair of LSL parents with differences in fruit firmness and anthocyanin concentration (05C197P002 × 16C108P065). These full-sib families were phenotyped for resistance to gray mold using an artificial inoculation protocol and genotyped with a 50K Axiom SNP array (Hardigan *et al*. 2020).

**Figure 2.**
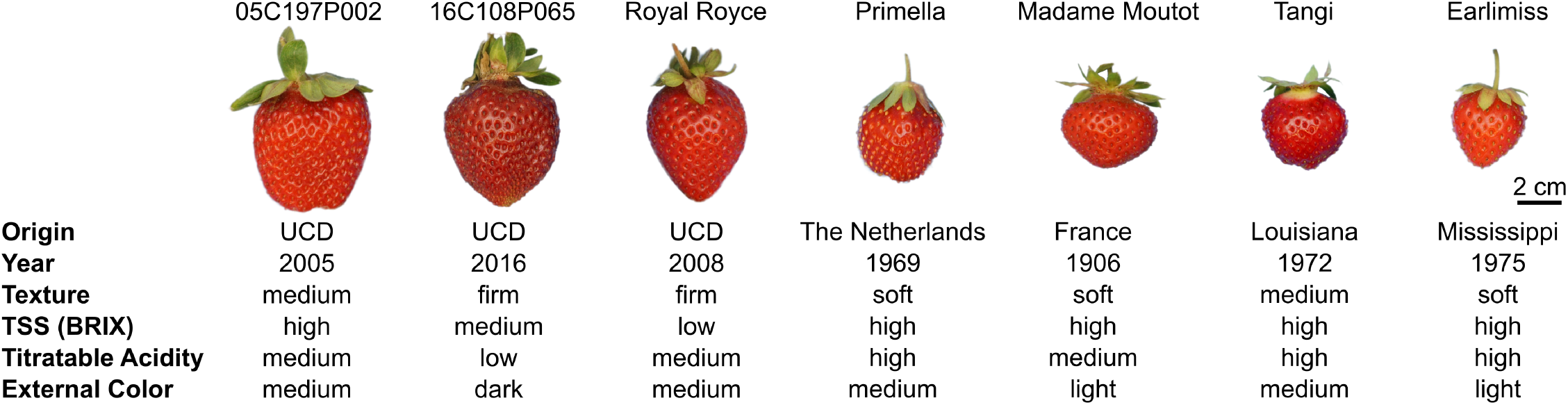
Fruit Phenotypes for Training Population Parents. Fruit of the parents of Royal Royce × Primella, Royal Royce × Madame Moutot, Royal Royce × Tangi, Royal Royce × Earlimiss, and 05C197P002 × 16C108P065 full-sib families. The countries and years of origin are shown for each parent. The fruit firmness categories were soft (< 0.15 kg/cm^2^), medium (0.15-0.30 kg/cm^2^), and firm (> 0.30 kg/cm^2^). The total soluble solids (TSS) categories were low (<9.0%), medium (9.0-11.0%), and high (>11.0%). Titratable acid (TA) concentration (%) categories were low (<0.7%), medium (0.7-1.0%), and high (>1.0%). External color intensity (L) categories were light (L >41.0), medium (25.0-40.0 L), and dark (<25.0 L).

### Development of a Highly Repeatable Protocol for Gray Mold Resistance Phenotyping in Strawberry

Natural infections are too inconsistent and unreliable for analyses of the genetics of resistance to gray mold in strawberry. To overcome this problem, we developed a highly repeatable artificial inoculation protocol for gray mold resistance phenotyping that involved puncturing fruit with a 3 mm probe, propagating spores of a single *B. cinerea* strain (B05.10), introducing a known concentration of spores into the wound site, and monitoring disease development on individual fruit stored undisturbed under high humidity (Fig. 3). Two quantitative *B. cinerea* disease symptoms were recorded on multiple fruits harvested from training population individuals: water-soaked lesion diameter (LD) in mm and the number of days post-inoculation (dpi) when external mycelium was observed on the surface of the fruit (EM). We found that incubating artificially inoculated fruit at 10°C and 95% humidity in the dark yielded highly repeatable results with minimal contamination from other postharvest decay pathogens. LD and EM were recorded daily from 1 to 14 dpi (Fig. 3-4). This protocol produced highly reproducible results with repeatability estimates in the 0.66-0.83 range for LD and 0.68-0.71 range for EM (Table 1). Although critical for maximizing repeatability, this protocol produced more severe disease symptoms than those commonly observed from natural infection, especially on non-wounded fruit of firm-fruited LSL cultivars (Jarvis 1962; Petrasch *et al*. 2019a) (Fig. 1 and 3; Fig. S1).

**Figure 3.**
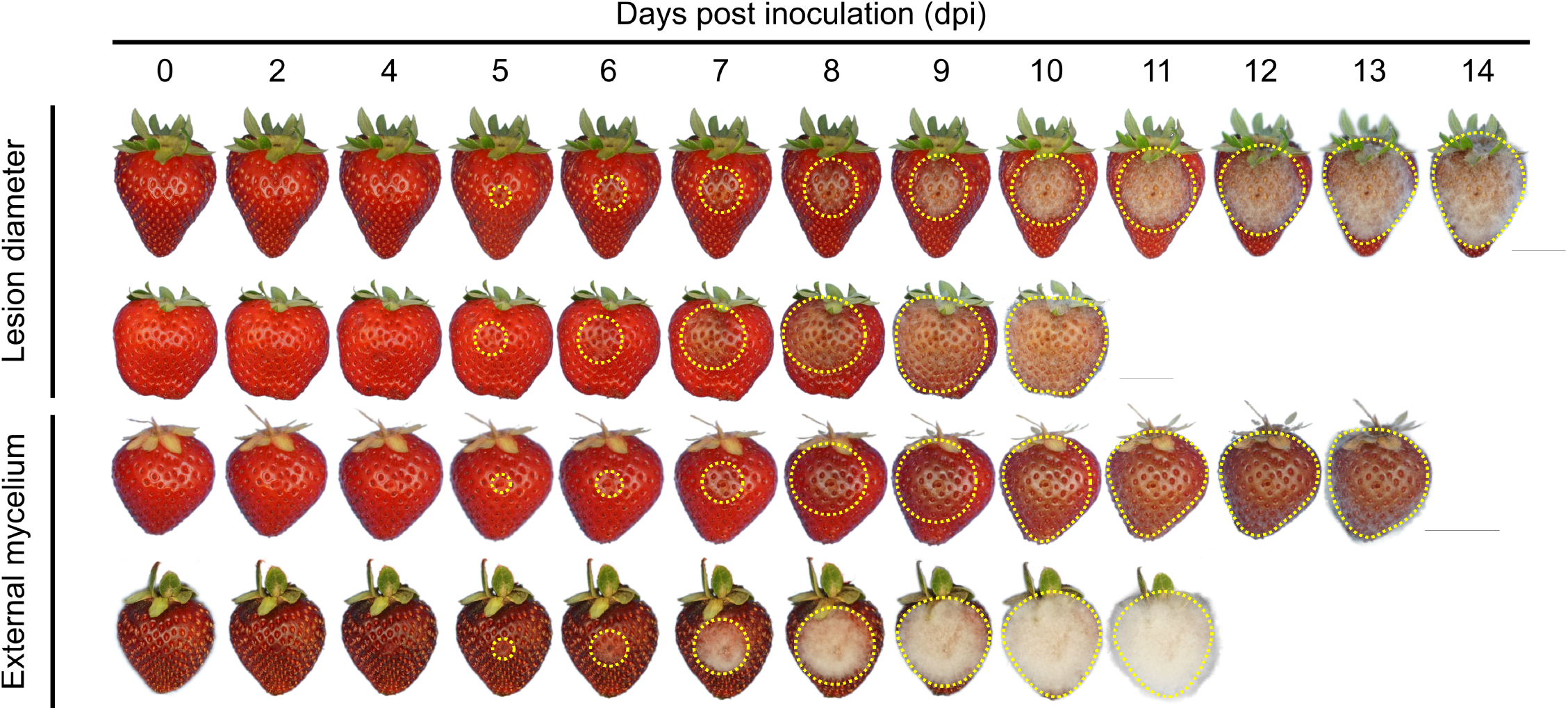
Postharvest Progression of Gray Mold Disease. Fruit were artificially inoculated with the *B. cinerea* strain B05.10 and phenotyped for 0 to 14 days post-inoculation (dpi). The dotted yellow lines highlight approximate lesion boundaries. The upper panel displays exemplary phenotypic extremes for lesion diameter (cm) among individuals in the multi-family training population. The 18C346P032 individual (first row) had one of the smallest lesion diameters at 8 dpi, whereas the 18C346P030 individual (second row) had one of the largest lesion diameters at 8 dpi. The lower panel displays exemplary phenotypic extremes for the speed of appearance of external mycelium on the fruit surface (dpi). The 18C346P025 individual (third row) had one of the slowest, whereas the 18B168P085 individual (fourth row) had one of the fastest times to the appearance of mycelium on the surface of the fruit.

**Table 1.** Repeatability (*r*) and Narrow-Sense Genomic Heritability (*h*^2^) Estimates for Gray Mold Lesion Diameter (LD) and Speed of Emergence of External Mycelium (EM)

| Population | Trait | $\hat{r}$ | $\hat{h}^2$ | |
| --- | --- | --- | --- | --- |
|  |  |  | Complete Subsamples | Single Subsample |
| Multi-Family | LD | 0.66 | 0.38 | 0.13 |
|  | EM | 0.68 | 0.39 | 0.16 |
| Royal Royce × Tangi | LD | 0.83 | 0.71 | 0.32 |
|  | EM | 0.71 | 0.44 | 0.13 |

Statistics were estimated for $n = 380$ individuals and $s = 1,520$ subsamples in a multi-family population and $n = 233$ individuals and $s = 1,386$ subsamples in the Royal Royce × Tangi population. The full-sib families in the multi-family population were Royal Royce × Earlimiss, Royal Royce × Madame Moutot, Royal Royce × Primella, Royal Royce × Tangi, and 05C197P002 × 16C108P06. Narrow-sense genomic heritability was estimated for 100% of the subsamples ( $\bar{s} = 2.9$ ) and for a single randomly selected subsample/individual ( $\bar{s} = 1.0$ ) in both populations, where $\bar{s}$ is the harmonic mean number of subsamples/individual.

### Genetics of Resistance to Gray Mold in Strawberry

To study the genetics of resistance to gray mold in strawberry, artificially inoculated fruit of individuals in multi-family and Royal Royce × Tangi populations were phenotyped daily for LD and EM over 14 days in cold storage (Fig. 3). The speed of fungal development and symptom severity differed among individuals in both populations (Fig. 4). The phenotypic extremes we observed are illustrated in time-series photographs of four individuals from the upper and lower tails of the LD and EM distributions in the multi-family population (Fig. 3). Lesions became visible and had enlarged to 10.0 mm by 5 dpi in one of the most susceptible individuals (18C346P030), whereas lesions were not visible until 8 dpi and developed the slowest in one of the least susceptible individuals (18C346P032). Lesions spanned the entire fruit surface of the most susceptible individuals by 8 dpi, thereby resulting in significant postharvest fruit deterioration and fungal decay (Fig. 3). Consequently, our genetic analyses of lesion diameter were applied to phenotypes observed 8 dpi, the last day in the study that resistance phenotypes could be observed for every individual.

**Figure 4.**
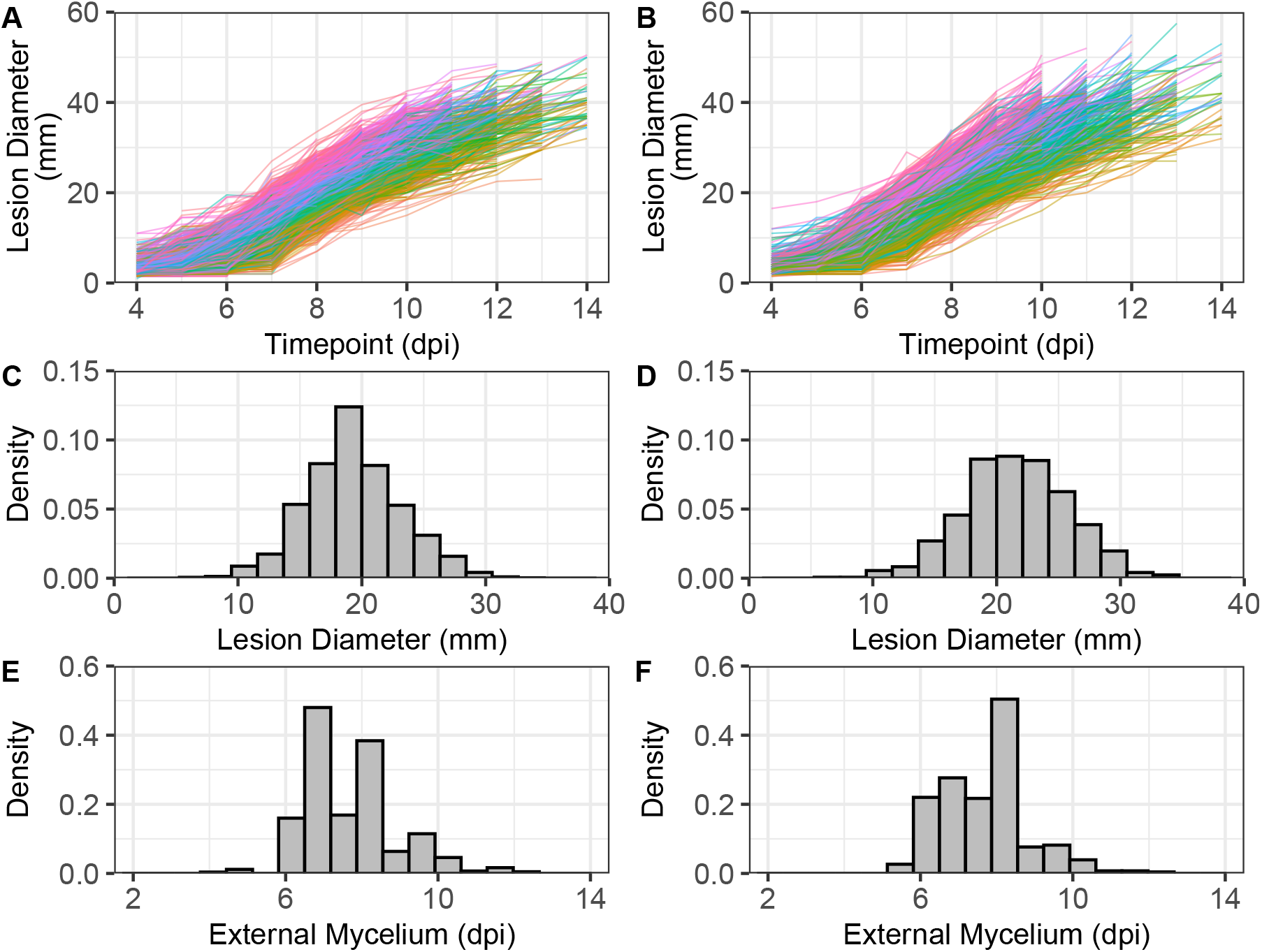
Distributions for Gray Mold Resistance Phenotypes. Fruit of individuals from a multi-family population (*n* = 380 individuals and *s* = 1,520 fruit) and the Royal Royce × Tangi population (*n* = 233 individuals and *s* = 1,386 fruit) were artificially inoculated with *B. cinerea* and phenotyped 0 to 14 days post-inoculation (dpi) for lesion diameter (LD; mm) and the speed of emergence of external mycelium on the surface of the fruit (EM; dpi). The multi-family population was grown in 2018-19, whereas independent samples of Royal Royce *times* Tangi individuals were grown in 2018-19 and 2019-20. (A) Estimated marginal means (EMM) for 380 individuals in the multi-family population were estimated for each time point from 11,163 observations (1,520 fruit × 4-14 time points) and plotted to visualize changes in LD over time. Because fruit periodically perished and were discarded, the number of observations was less than 21,280 (1,520 fruit × 14 time points). The curves for each individual were plotted with colors corresponding to the EMM rank for lesion diameter at 8 dpi. (B) EMMs for 233 individuals in the Royal Royce × Tangi population were estimated from 9,797 observations (1,386 fruit × 4-14 time points) and plotted and color highlighted as described for the multi-family population. (C and E) Phenotypic distributions for LD and EM in the multi-family population. (D and F) Phenotypic distributions for LD and EM in the Royal Royce × Tangi population.

As expected, our analyses confirmed that resistance to gray mold is genetically complex in strawberry, a finding consistent with observations in other hosts (Lurie *et al*. 1997; Glazebrook 2005; Rowe and Kliebenstein 2008; Lewers *et al*. 2012; Corwin *et al*. 2016; Hanson *et al*. 2016; Petrasch *et al*. 2019a; Zhang *et al*. 2019; Silva *et al*. 2020; Caseys *et al*. 2021). Although statistically significant differences were observed among individuals for LD and EM in both populations (*p* < 0.01), every individual was susceptible and the phenotypic ranges were comparatively narrow (Fig. 3-4). Lesion diameters were approximately normally distributed and ranged from 7.0 to 33.5 mm at 8 dpi in the multi-family population and 7.0 to 34.0 mm at 8 dpi in the Royal Royce × Tangi population (Fig. 4; Fig. S2). Similarly, the speed of appearance of mycelium on the surface of the fruit (EM) was approximately normally distributed and ranged from 4.0 to 12.5 dpi in the multi-family population and 5.5 to 12.5 dpi in the Royal Royce × Tangi population (fruit were phenotyped out to 14 dpi). The repeatabilities for LD and EM among individuals in these populations suggested that one-third or more of the phenotypic variation observed for gray mold resistance was genetically caused (Table 1). Narrow-sense genomic heritability estimates ranged from 0.38 to 0.71 for LD and 0.39 to 0.44 for EM, which suggested that a significant fraction of the genetic variation was additive and thus that resistance to gray mold can be enhanced by artificial selection (Table 1).

Genome-wide searches failed to identify large-effect loci for LD or EM (Fig. 5; Table 2). These searches included GWAS in the training populations and QTL mapping in individual full-sib families. Nine family-specific QTL with small effects were identified, eight for LD and one for EM (Table 2). None had effects large enough to warrant targeting by marker-assisted selection or inclusion as fixed effects in genomic prediction models. Although the QTL effects were small and family-specific (Table 2), a few interesting candidate gene associations were identified when short QTL-associated haploblocks were searched in the reference genome for genes with biotic stress and disease resistance annotations (File S5). A cluster of 11 tandemly duplicated genes encoding pathogenesis-related (PR) proteins were found in close proximity to the most significant SNP (AX-184469645) associated with a QTL on chromosome 4A (Table 2). These genes share sequence homology to *FcPR10*, an *F. chiloensis* ribonuclease encoding gene previously predicted to reduce the severity of gray mold disease in strawberry (González *et al*. 2009, 2013). The other QTL-associated candidate genes that might warrant further study encode peroxidases (chromosome 7D; Mb 576-2,709) reported to modulate reactive oxygen species levels and inhibit fungal growth during *B. cinerea* infections (Cantu *et al*. 2008, 2009; Tomas-Grau *et al*. 2018) and transcription factors reported to signal pathogen-triggered immunity, e.g., WRKY and AP2/ERF (chromosome 3B; Mb 13,870-15,400) (Gutterson and Reuber 2004; Bigeard *et al*. 2015), that might target pathogenicity factors, e.g., chitinases (van Schie and Takken 2014) and protease inhibitors (Hermosa *et al*. 2006; Billon-Grand *et al*. 2012). While these genes are worthwhile candidates for further study (González *et al*. 2009, 2013; Petrasch *et al*. 2019a), the effects of the associated QTL were too small and insignificant for direct selection (Table 2). This was, nevertheless, a first attempt to identify loci underlying resistance to *B. cinerea* in strawberry through a genome-wide search for genotype-to-phenotype associations in the octoploid genome (File S5).

**Figure 5.**
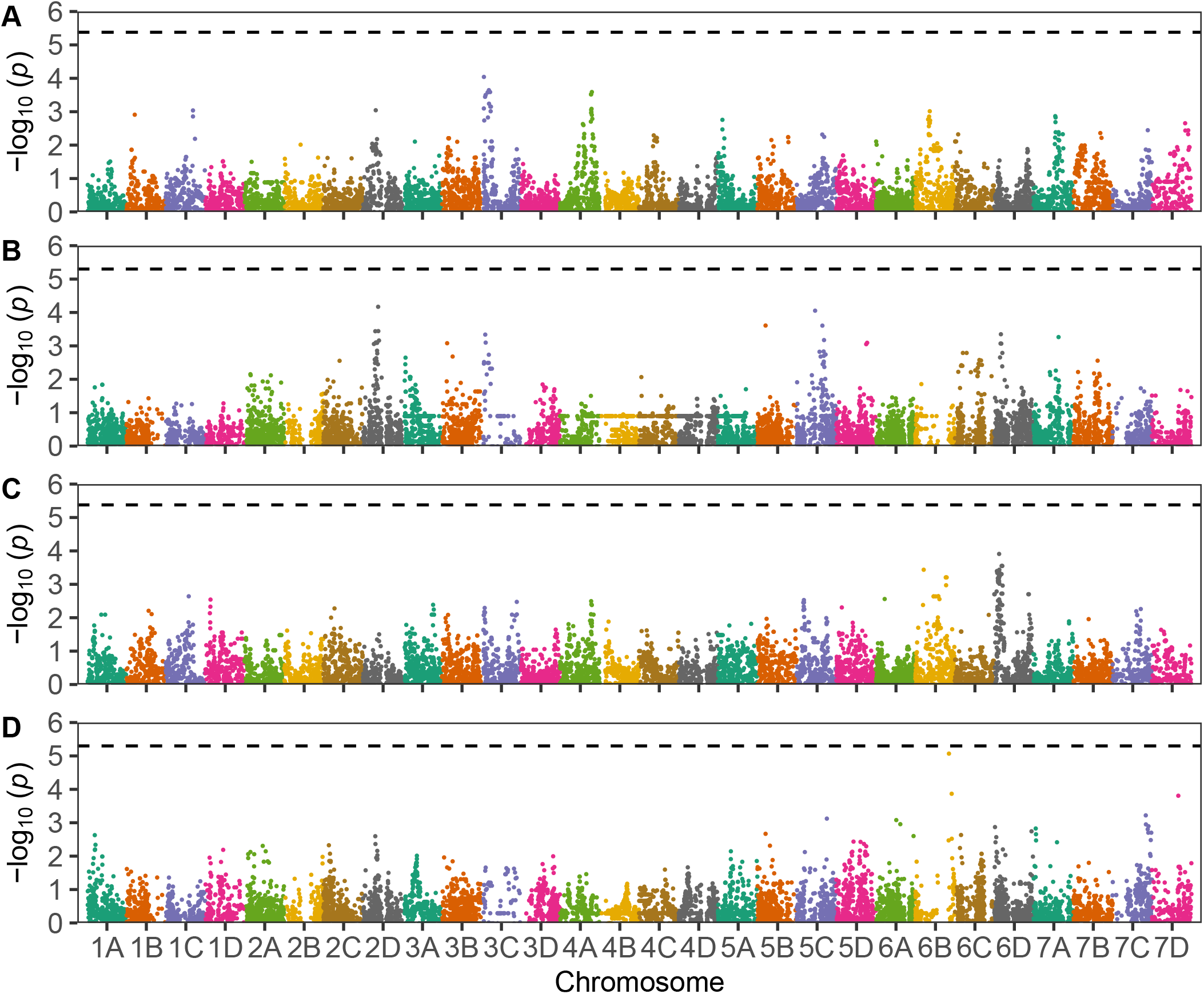
Genome-Wide Association Study of Gray Mold Resistance. Manhattan plots are shown for genome-wide scans for marker-trait associations for lesion diameter (LD) at 8 days post-inoculation (dpi) in the multi-family (A), LD at 8 dpi in the Royal Royce × Tangi (B), speed of emergence of external mycelium on the surface of the fruit (EM) in the multi-family (C), and EM in the Royal Royce × Tangi (D) populations. The individuals in both populations were genotyped with a 50K Axiom SNP array. The analyses were done using the ‘Camarosa’ reference genome (Edger *et al*. 2019) with physical positions of SNP markers ascertained by (Hardigan *et al*. 2020) using the chromosome nomenclature of (Hardigan *et al*. 2021). Horizontal dashed lines identify the genome-wide Bonferroni significance threshold.

**Table 2.** Summary Statistics for Quantitative Trait Loci Affecting Gray Mold Resistance^1^ in Strawberry.

| Population | SNP Marker<sup>2</sup> | Chromosome<sup>3</sup> | Position (Mb)<sup>4</sup> | Contrast<sup>5</sup> | Effect | LOD<sup>6</sup> | Allele<sup>7</sup> |
| --- | --- | --- | --- | --- | --- | --- | --- |
| Lesion Diameter (mm) |  |  |  |  |  |  |  |
| 05C197P002 × 16C108P065 | AX-184345814 | 7D | 0.0 | AA-AB | -3.48 | 4.03 | 05C197P002 |
| Royal Royce × Primella | AX-184496623 | 3B | 105.0 | BB-AB | 2.49 | 3.61 | Royal Royce |
| Royal Royce × Primella | AX-184685020 | 5C | 88.7 | AA-AB | -0.87 | 3.70 | Royal Royce |
| Royal Royce × Earlimiss | AX-184026159 | 3C | 24.2 | AA-AB | -2.79 | 5.36 | Royal Royce |
| Royal Royce × Earlimiss | AX-184469645 | 4A | 101.6 | AA-AB | 2.52 | 4.26 | Earlimiss |
| Royal Royce × Tangi | AX-184718804 | 3C | 52.8 | BB-AB | 1.92 | 4.21 | Royal Royce |
| Royal Royce × Tangi | AX-184031508 | 5A | 0.0 | BB-AB | 1.92 | 4.56 | Royal Royce |
| Royal Royce × Tangi | AX-184266150 | 7C | 21.7 | AA-AB | 1.82 | 3.70 | Tangi |
| External Mycelium (dpi) |  |  |  |  |  |  |  |
| Royal Royce × Tangi | AX-184857528 | 5C | 93.1 | BB-AB | 0.48 | 4.03 | Tangi |

<sup>1</sup>Lesion diameter (LD) and the speed of emergence of external mycelium on the surface of the fruit (EM) were recorded daily from 0 to 14 days post-inoculation (dpi). QTL statistics are shown for LD at 8 dpi, the last day that none of the fruit had perished.
<sup>2</sup>Alphanumeric names for SNP markers on the 50K Axiom SNP array Hardigan et al. \(2020\) with the largest effects (largest LOD score) for a particular QTL.
<sup>3</sup>Chromosome numbers follow the nomenclature proposed by Hardigan et al. \(2021\) with letters designating subgenomes (A-D) and numbers designating homeologous chromosomes (1-7).
<sup>4</sup>The physical positions of SNP markers were ascertained by (Hardigan et al. 2020) from alignments of SNP probe sequences to the *F. × ananassa* 'Camarosa' reference genome (Edger et al. 2019).
<sup>5</sup>SNP marker genotypes were coded AA, AB, and BB where A is the allele transmitted by the parent shown on the left in the pedigree and B is the allele transmitted by the parent shown on the right in the pedigree. Contrasts were estimated for each SNP marker as the difference between estimated marginal means (EMMs) for genotypes. The AA-AB contrast compared EMMs between AA homozygotes and AB heterozygotes, whereas the BB-AB contrast compared EMMs between BB homozygotes and AB heterozygotes.
<sup>6</sup>Logarithm of the odds (LOD) scores for SNP marker-QTL associations that exceeded statistical thresholds estimated by permutation.
<sup>7</sup>The parent that transmitted the favorable allele. The favorable allele for LD decreased the EMM, whereas the favorable allele for EM increased the EMM.

Royal Royce, the firm-fruited LSL parent, was more resistant to gray mold than the soft-fruited SSL parents (phenotypic distributions and parent EMMs for each full-sib family are shown in Fig. S2). Lesions were smaller and mycelium appeared later in Royal Royce than the other parents (resistance increased as LD decreased and EM increased). Royal Royce was the more resistant parent for both traits in the four full-sib families with that parent (Fig. 2; Fig. S2). For the 05C197P002 × 16C108P065 full-sib family, 05C197P002 was more resistant than 16C108P065 for LD and vice versa for EM. The LD and EM differences were highly significant (p ≤ 0.01) with individuals transgressing the phenotypic ranges of the parents (Fig. S2). Transgressive segregation was primarily bidirectional for both traits; however, the EM distributions for Royal Royce × Tangi and 05C197P002 × 16C108P065 were right-skewed towards more resistance (slower external mycelium emergence) and lacked individuals in the lower tails distal to the more susceptible parent (Fig. S2). These results suggested that favorable alleles were transmitted by both parents for both traits and that favorable alleles for different loci segregated in most of the families.

Lesion diameters were plotted for every individual to visualize phenotypic changes in disease symptoms over time (Fig. 4A-B). The heatmap colors of the individual curves were determined from the lesion diameter EMM ranks at 8 dpi. These plots show that the speed of lesion development differed among individuals, that cross-over individual × time interactions were negligible, and that the phenotypic changes among individuals were approximately parallel over time, all of which increased confidence in the heritability of the phenotypic differences we observed (Table 1).

### Genomic Prediction Accuracies Varied Between Populations and Symptoms

Genomic prediction accuracies for different whole-genome regression (WGR) methods ranged from 0.28 to 0.47 for LD and 0.37 to 0.59 for EM when estimated by cross-validation from 100% of the subsamples (Table 3; Fig. 6-7A, D, G, and J; Fig. S3). The differences in accuracy among WGR methods were mostly negligible (Table 3; Fig. 6-7; Fig. S3). The prediction accuracy was greater for LD than EM in the Royal Royce × Tangi population (Fig. 6G and J), whereas the reverse was observed in the multi-family population (Fig. 6A and D). Using cross-validation with 100% of the subsamples, clear differences in prediction accuracy and shrinkage were observed between disease symptoms within and between populations (Table 3; Fig. 6-7A, D, G, and J); Fig. S3). The prediction accuracy for LD was markedly different between the multi-family and Royal Royce × Tangi populations (Fig. 6A and C). The GEBV range for LD in the multi-family population was half as wide (15.5 to 22.8) and the kernel density was flatter and more vertical than that observed in the Royal Royce × Tangi population (13.3 to 27.4) (Fig. 6A and G). Notably, the LD phenotypes of the most resistant individuals in the RR × Tangi population (those with the smallest LD means) were well predicted (Fig. 6G). Their EM phenotypes, however, were not as well predicted—the GEBV range for EM (6.8-9.2) was half that of the phenotypic range (5.5-10.7) and the kernel density distribution was flatter and more vertical (Fig. 6-7J; Fig. S3).

**Table 3.** Genomic Prediction Accuracy for Gray Mold Resistance.

| Genomic Prediction Accuracy |  |  |  |  |
| --- | --- | --- | --- | --- |
| Population<sup>1</sup> | WGR Method<sup>2</sup> | Complete Subsample Cross-Validation<sup>3</sup> | Single Subsample Cross-Validation<sup>4</sup> | Single Subsample GEBV Prediction<sup>5</sup> |
| Lesion Diameter (mm) |  |  |  |  |
| Multi-Family | G-BLUP | 0.33 | 0.17 | 0.39 |
|  | RKHS | 0.33 | 0.19 | 0.56 |
|  | SVM | 0.28 | 0.17 | 0.40 |
| Royal Royce × Tangi | GBLUP | 0.52 | 0.56 | 0.61 |
|  | RKHS | 0.59 | 0.57 | 0.71 |
|  | SVM | 0.59 | 0.58 | 0.63 |
| External Mycelium (dpi) |  |  |  |  |
| Multi-Family | G-BLUP | 0.47 | 0.35 | 0.44 |
|  | RKHS | 0.47 | 0.34 | 0.48 |
|  | SVM | 0.44 | 0.40 | 0.49 |
| Royal Royce × Tangi | G-BLUP | 0.37 | 0.36 | 0.43 |
|  | RKHS | 0.42 | 0.34 | 0.64 |
|  | SVM | 0.40 | 0.35 | 0.47 |

<sup>1</sup>Statistics were estimated from analyses of $n = 380$ individuals and $s = 1,520$ subsamples in a multi-family population and $n = 233$ individuals and $s = 1,386$ subsamples in the Royal Royce × Tangi population. The full-sib families in the multi-family population were Royal Royce × Earlimiss, Royal Royce × Madame Moutot, Royal Royce × Primella, Royal Royce × Tangi, and 05C197P002 × 16C108P06.
<sup>2</sup>Genomic estimated breeding values (GEBVs) were estimated using genomic-best linear unbiased prediction (G-BLUP), reproducing kernel Hilbert space (RKHS) regression, and support-vector machine (SVM).
<sup>3</sup>GEBVs and genomic prediction accuracies were estimated from 100% of the subsamples/individual by cross-validation with 1,000 permutations using 80% of the individuals for training and 20% of the individuals for prediction, where the harmonic mean number of subsamples/individual ( $\bar{s}$ ) = 2.9.
<sup>4</sup>Genomic prediction accuracy was estimated from a single randomly selected subsample/individual by cross-validation with 1,000 permutations using 80% of the individuals for training and 20% of the individuals for prediction.
<sup>5</sup>Correlation between GEBVs estimated from a single randomly selected subsample/individual and EMMs of individuals estimated from 100% of the subsamples/individual.

**Figure 6.**
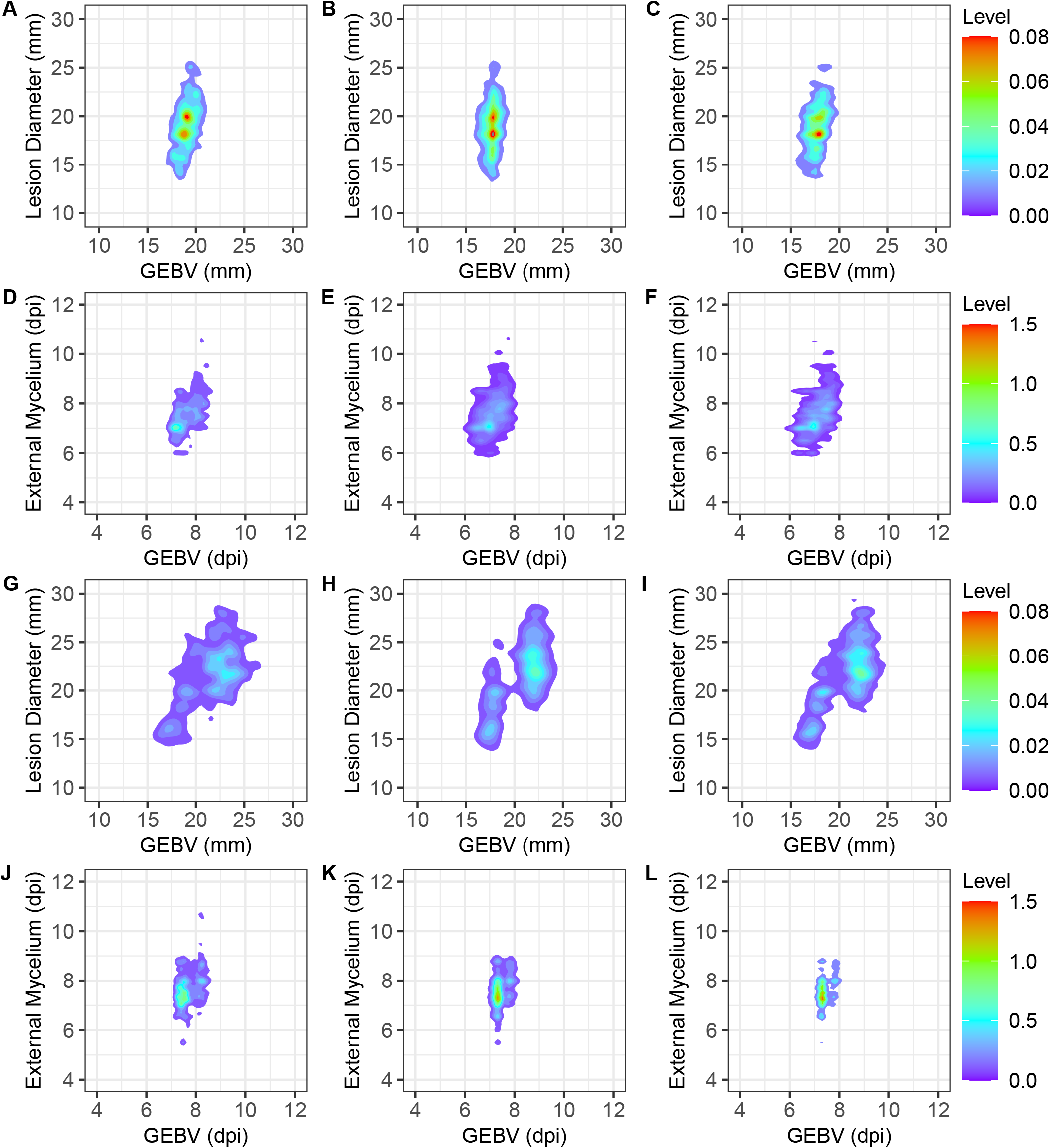
Cross-Validation Accuracy of Genomic Predictions for Gray Mold Resistance. (A-C) Phenotype by genomic estimated breeding value (GEBV) density plots are shown for LD and EM among individuals in a multi-family training population (A-F) and the Royal Royce × Tangi training population (G-L). GEBVs were estimated using G-BLUP. The density plots shown in the first column (A, D, G, and J) display estimates from 80:20 cross-validation using marginal means (EMMs) estimated from 100% of the subsamples/individual, where data for 80% of the individuals were used to estimate GEBVs for the other 20% of the individuals. There were 380 individuals and a mean of 2.4 subsamples/individual in the multi-family population (1,393 fruit were phenotyped daily) and 233 individuals and a mean of 3.6 subsamples/individual in the Royal Royce × Tangi population (1,373 fruit were phenotyped daily). The density plots shown in the middle column display estimates from 80:20 cross-validation using phenotypic observations for one randomly chosen subsample/individual, where data for 80% of the individuals were used to estimate GEBVs for the other 20% of the individuals. The density plots shown in the last column (C, F, I, and L) display GEBVs estimated from one randomly chosen subsample/individual and EMMs of those individuals estimated from 100% of the subsamples.

**Figure 7.**
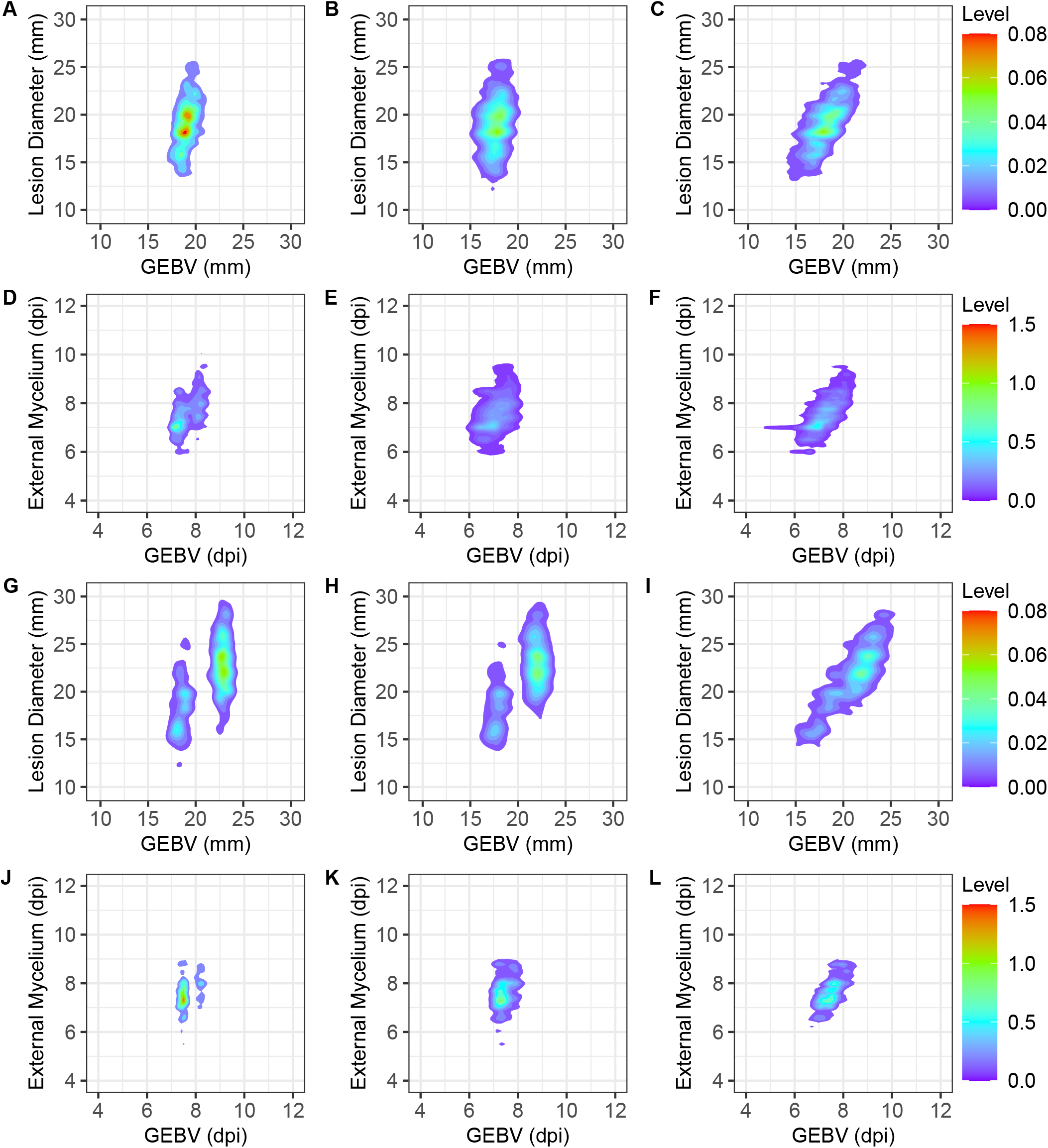
Cross-Validation Accuracy of Genomic Predictions for Gray Mold Resistance. (A-C) Phenotype by genomic estimated breeding value (GEBV) density plots are shown for LD and EM among individuals in a multi-family training population (A-F) and the Royal Royce × Tangi training population (G-L). GEBVs were estimated using reproducing kernel Hilbert spaces (RKHS) regression. The density plots shown in the first column (A, D, G, and J) display estimates from 80:20 cross-validation using marginal means (EMMs) estimated from 100% of the subsamples/individual, where data for 80% of the individuals were used to estimate GEBVs for the other 20% of the individuals. There were 380 individuals and a mean of 2.4 subsamples/individual in the multi-family population (1,393 fruit were phenotyped daily) and 233 individuals and a mean of 3.6 subsamples/individual in the Royal Royce × Tangi population (1,373 fruit were phenotyped daily). The density plots shown in the middle column display estimates from 80:20 cross-validation using phenotypic observations for one randomly chosen subsample/individual, where data for 80% of the individuals were used to estimate GEBVs for the other 20% of the individuals. The density plots shown in the last column (C, F, I, and L) display GEBVs estimated from one randomly chosen subsample/individual and EMMs of those individuals estimated from 100% of the subsamples.

One of the challenges of breeding for resistance to gray mold and other postharvest traits is phenotyping throughput. Collectively, 2,563 fruit were harvested and individually stored, tracked, and phenotyped in our study (Fig. 4). Our expectation was that multiple fruit/individual were needed to more accurately estimate EMMs and GEBVs and nominally increase heritability. To assess the effect of subsamples on prediction accuracy and explore the feasibility of applying selection for resistance to gray mold from a single subsample/individual, GEBVs and prediction accuracies were estimated from a single randomly selected subsample/individual. We observed a significant decrease in narrow-sense genomic heritability for LD and EM in the single subsample analyses, e.g., *ĥ*^2^ decreased from 0.38 to 0.13 for LD and 0.39 to 0.16 for EM in the multi-family population (Table 1). Naturally, prediction accuracies plummeted in the single subsample analyses too (Table 3; Fig. 6-7 and Fig. S3). This is clearly illustrated by the kernel density distributions for genomic selection accuracy estimated for G-BLUP, RKHS, and SVM by cross-validation with a single sub-sample/individual (Fig. 6-7B, E, H, and K; Fig. S3). GEBV ranges were narrower and kernel density distributions were flatter and more vertical for the single subsample versus multiple subsample analyses for LD and EM in both populations (Fig. 6-7; Fig. S3). Hence, we concluded that breeding values cannot be accurately predicted without subsampling fruit. Nevertheless, a shortcoming of this study was that genomic prediction accuracies could not be compared for equivalent numbers of phenotypic observations for individuals with and without subsamples.

To shed more light on the effect of subsamples, GEBVs were estimated from a single subsample/individual from 100% of the individuals in each population (Table 3; Fig. 6C, F, I, and L). This analysis was less stringent than either of the cross-validation analyses because GEBV estimates were directly compared to EMMs. The prediction accuracies for these analyses were slightly greater than those estimated by 80:20 cross-validation (Table 3; Fig. 6-7 C, F, I, and L). These results, while less stringent than the cross-validation analyses, suggested that increasing the number of individuals with fewer subsamples/individual might yield predictions as accurate as those achieved with fewer individuals and an increased number of subsamples/individual, although this needs to be empirically tested and validated.

### Gray Mold Resistance Traits Were Genetically Correlated With Shelf Life-Associated Fruit Quality Traits

One of our original hypotheses was that selection for increased fruit firmness and other shelf life-associated fruit quality traits pleiotropically decreased susceptibility to gray mold in strawberry. The additive genetic correlations observed among the traits pheno typed in the present study support this hypothesis (Fig. 8). LD and EM were weakly negatively genetically correlated 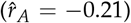. These gray mold resistance traits were weakly to strongly correlated with fruit quality traits in directions predicted by our hypotheses. Because gray mold resistance increases as LD decreases and EM increases, signs of the additive genetic correlations have different interpretations for LD and EM and can be antagonistic or synergistic. The interpretation depends on the specific phenotypes targeted for a particular market, e.g., SSL versus LSL.

**Figure 8.**
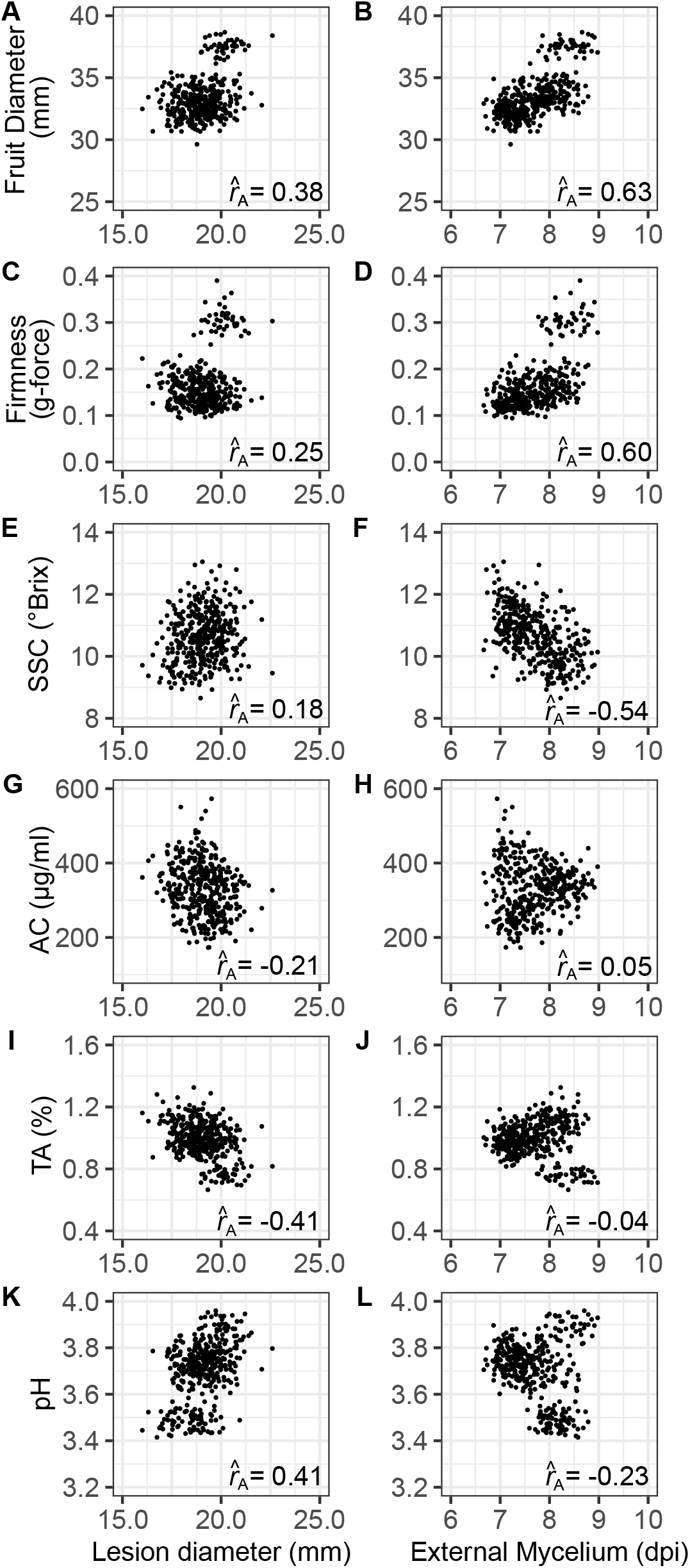
Additive Genetic Correlations Between Gray Mold Resistance and Fruit Quality Traits. Scatter plots are shown for G-BLUP estimates of genomic-estimated breeding values (GEBVs) for gray mold resistance and fruit quality traits among 380 individuals in a multi-family training population. Additive genetic correlations 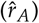 were estimated between lesion diameter (LD), the speed of emergence of external mycelium on the surface of the fruit (EM), fruit diameter, firmness, soluble solids content (SSC), anthocyanin content (AC), titratable acidity (TA), and pH.

LD was negatively genetic correlated with titratable acidity 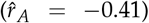 and positively genetically correlated with pH 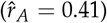; hence, LD increased as titratable acidity decreased and pH increased (Fig. 8I-J). The effect of titratable acidity on resistance phenotypes was the motivation for screening additional individuals from the Royal Royce × Tangi family, which had significant genetic variation for TA and yielded more accurate genomic predictions for LD than were observed in the multi-family population (Fig. 6-7; Fig. S3). EM was more strongly positively genetically correlated with fruit size and firmness than LD and negatively genetically correlated with total soluble solids (°BRIX) 8. EM increased (disease resistance increased) as BRIX decreased and fruit size and firmness increased. Hence, we found that mycelium developed faster on softer, sweeter, and smaller fruit than firmer, less sweet, and larger fruit, as is typical of Royal Royce and other modern LSL cultivars (Fig. 1-).

Finally, LD was weakly negatively genetically correlated with anthocyanin concentration, whereas EM was uncorrelated with anthocyanin concentration (Fig. 8G-H). Although the additive genetic correlation we observed between LD and anthocyanin concentration was in the direction predicted by previous studies in strawberry and tomato (Jersch *et al*. 1989; Hébert *et al*. 2002; Bassolino *et al*. 2013; Zhang *et al*. 2013), lesion diameter only slightly decreased as anthocyanin concentration increased (Fig. 8).

### Breeding for Enhanced Resistance to Gray Mold in Strawberry

Breeding for resistance to necrotrophic pathogens has been challenging in plants (Finkers *et al*. 2007b,a; Williamson *et al*. 2007; Petrasch *et al*. 2019a; Delplace *et al*. 2020; Lorang 2019). The mechanisms of resistance to necrotrophic pathogens are more subtle, quantitative, and complex than those commonly observed for biotrophic pathogens that trigger pathogen-associated molecular pattern-triggered-immunity and effector-triggered immunity (Glazebrook 2005; Jones and Dangl 2006; Jones *et al*. 2016; Saijo *et al*. 2018; Zhang *et al*. 2019; Lorang 2019; Caseys *et al*. 2021). Our findings were well aligned with previous findings in other *B. cinerea* hosts and shed light on the genetic complexity of resistance to gray mold in strawberry. Where do we go from here? We are skeptical that significant genetic gains can be achieved for gray mold resistance across the complete shelf life spectrum in strawberry but are confident that postharvest gray mold incidence can be minimized but obviously not eliminated in long shelf life populations. This conclusion seems well aligned with previous findings in strawberry and other hosts of this pathogen (Finkers *et al*. 2007b,a; Rowe and Kliebenstein 2008; Lewers *et al*. 2012). Because several fruit quality traits pleiotropically affect gray mold resistance in strawberry, the challenge is exponentially greater when breeding for markets where softer fruits with elevated sugars are preferred and long shelf life phenotypes are neither necessary nor preferred (Fig. 1-2 and 8). However, for markets where LSL cultivars are essential, direct selection for the requisite fruit quality traits seems to confer sufficient resistance to gray mold to ensure marketability under normal postharvest storage conditions and timelines, especially for fruit produced in coastal California and other arid and semi-arid environments with low humidity and rainfall (Fig. 1; Fig. S1). Although we only sampled 12 coastal California environments (six locations × two harvests/location) to estimate the natural incidence of *B. cinerea* among eight LSL cultivars, we suspect that deeper sampling will confirm our findings.

There are open questions to be addressed and were limitations to our study. First, we did not screen diverse germplasm to identify sources of resistance to gray mold. As our study and others have shown, strong sources of resistance to this pathogen may not exist (Chandler *et al*. 2004; Seijo *et al*. 2008; Lewers *et al*. 2012; Bestfleisch *et al*. 2015). The narrow-sense heritability estimates and genomic prediction results for LD and EM in the present study suggest that a deeper exploration of genetic diversity, while challenging, seems worthwhile (Table 1; Fig. 6-7; Fig. S3). The association between resistance and titratable acidity seems to be particularly promising and worthy of further study (Fig. 8), particularly if increased acidity is offset by increased sugars to achieve a palatable sugar:acid balance.

Second, the parents for this study were selected to assess the effects of fruit quality and shelf life-associated traits on gray mold disease development (Fig. 2), not for *known* intrinsic differences in gray mold resistance that are genetically uncorrelated with fruit quality and shelf life phenotypes. The pleiotropic effects of shelf life-associated fruit quality traits on gray mold susceptibility appear to be inescapable (Fig. 8). Our results suggest that resistance can be increased by selecting for increased titratable acidity and firmness and decreased sugars but these phenotypes profoundly affect flavor and cannot be manipulated in a vacuum (Zorrilla-Fontanesi *et al*. 2011; Diamanti *et al*. 2012; Lerceteau-Köhler *et al*. 2012; Verma *et al*. 2017). Without more extensive germplasm screening, the data needed to guide the selection of parents for future genetic studies are lacking.

Third, the natural incidence of gray mold was only explored in the present study for LSL cultivars commercially grown in California (Fig. 1). Highly perishable short- and medium-shelf life cultivars, as typified by the heirloom cultivars (parents) we screened (Fig. 2), are challenging to grow and phenotype in such studies because they typically have low yields, are easily bruised and wounded, and cannot be harvested and handled with the same robustness as commercially important LSL cultivars. Nevertheless, a study of the natural incidence of gray mold among individuals spanning the shelf life spectrum could shed further light on genetic correlations between fruit quality and gray mold resistance phenotypes and possibly identify sources of favorable alleles underlying intrinsic resistance that are uncorrelated with fruit quality traits, e.g., biochemical phenotypes triggered by defense mechanisms (Diaz *et al*. 2002; Glazebrook 2005; van Kan 2006; Williamson *et al*. 2007; Veloso and van Kan 2018; Lorang 2019; Petrasch *et al*. 2019a). Because gray mold disease symptoms from artificial inoculation protocols are typically harsher than those observed from natural infections of non-wounded fruit (Fig. 1 and 3), a deeper exploration of the natural incidence of gray mold seems warranted in strawberry, perhaps by simulating rainfall in field experiments through overhead irrigation or other practices to increase the uniformity and incidence of natural infection.

Our results suggest that phenotypic or genomic selection could be effective for gray mold resistance but only in certain populations and only when selection for genetically correlated traits does not antagonistically reduce genetic gains for gray mold resistance phenotypes (Table 3; Fig. 6-7; Fig. S3). Genetic gains for gray mold resistance are affected by shelf-life related traits through additive genetic correlations and could be reversed by simultaneous selection for fruit quality and shelf life traits that antagonistically pleiotropically affect gray mold resistance phenotypes (Fig. 8). Genetic variation for fruit quality traits strongly affected the phenotypic differences we observed for LD and EM. Most importantly, the fruit quality traits associated with enhanced flavor were antagonistically genetically correlated with gray mold resistance phenotypes (Fig. 8).

Cross-validation of genomic predictions in the present study shed light on the complexity of genetic mechanisms underlying gray mold resistance phentoypes and highlighted the challenges inherent in breeding for increased resistance to gray mold in strawberry (Table 3; Fig. 6-7; Fig. S3). The three WGR methods we applied to the prediction problem strongly shrunk breeding values to the population mean for LD in the multi-family and EM in the Royal Royce × Tangi populations. Such shrinkage is typical for moderately heritable complex diseases in plants (Rutkoski *et al*. 2011, 2014; Poland and Rutkoski 2016).

The prospects for identifying superior genotypes through genomic selection were greater for LD in the Royal Royce × Tangi population and EM in the multi-family population than vice versa. Whether applying phenotypic or genomic selection, the probability of selecting superior genotypes can be exceedingly low when breeding for resistance to genetically complex diseases in plants (Poland and Rutkoski 2016; Crossa *et al*. 2017; Voss-Fels *et al*. 2019). Nevertheless, with cost-effective genome-wide genotyping, genomic selection has the potential to increase genetic gains by increasing the number of selection candidates that can be screened per unit of time and space (Poland and Rutkoski 2016; Crossa *et al*. 2017). Even in the small populations we studied (*n* = 380 and *n* =233), nearly 20,000 phenotypic observations were collected to quantitatively assess resistance to gray mold, which included image analyses at each post-inoculation time point (Fig. 4). The phenotyping throughput needed to effectively evaluate postharvest traits in strawberry can be limiting, particularly when multiple harvests are factored into the equation, e.g., day-neutral cultivars are typically harvested twice weekly over a period of six to eight months in coastal California. Because the progression of disease symptoms was continuous and genotype × storage time interactions were negligible (Fig. 4), phenotyping throughput can be increased by employing a response surface experiment design, e.g., by phenotyping fruit at three equally spaced time points spanning the range needed to estimate response curves and accurately predict LD and EM phenotypes (Hill and Hunter 1966; Khuri and Mukhopadhyay 2010). This is precisely the sort of breeding problem where genomic prediction methods have the greatest potential utility and applicability (Heffner *et al*. 2010; Jannink *et al*. 2010; Lin *et al*. 2014), despite the complexity of the underlying genetic mechanisms and variable prediction accuracies across populations, which nevertheless affect phenotypic and genomic selection equally (Table 3; Fig. 6-7; Fig. S3).

## ACKNOWLEDGEMENTS

The authors thank Nancy N. Her and Ozalique Williams for assisting with disease assessments and Nayeli Valencia De Puglisi, Bruce Campopiano, and Eduardo Garcia for assisting with fruit harvests.

## CONFLICT OF INTEREST

The authors declare no conflict of interest.

